# Characterizing RNA Pseudouridylation by Convolutional Neural Networks

**DOI:** 10.1101/126979

**Authors:** Xuan He, Sai Zhang, Yanqing Zhang, Tao Jiang, Jianyang Zeng

## Abstract

The most prevalent post-transcriptional RNA modification, pseudouridine (Ψ), also known as the fifth ribonucleoside, is widespread in rRNAs, tRNAs, snRNAs, snoRNAs and mRNAs. Pseudouridines in RNAs are implicated in many aspects of post-transcriptional regulation, such as the maintenance of translation fidelity, control of RNA stability and stabilization of RNA structure. However, our understanding of the functions, mechanisms as well as precise distribution of pseudourdines (especially in mRNAs) still remains largely unclear. Though thousands of RNA pseudouridylation sites have been identified by high-throughput experimental techniques recently, the landscape of pseudouridines across the whole transcriptome has not yet been fully delineated. In this study, we present a highly effective model, called PULSE (<u>P</u>seudo<u>U</u>ridy<u>L</u>ation <u>S</u>ites <u>E</u>stimator), to predict novel Ψ sites from large-scale profiling data of pseudouridines and characterize the contextual sequence features of pseudouridylation. PULSE employs a deep learning framework, called convolutional neural network (CNN), which has been successfully and widely used for sequence pattern discovery in the literature. Our extensive validation tests demonstrated that PULSE can outperform conventional learning models and achieve high prediction accuracy, thus enabling us to further characterize the transcriptome-wide landscape of pseudouridine sites. Overall, PULSE can provide a useful tool to further investigate the functional roles of pseudouridylation in post-transcriptional regulation.

## Introduction

Pseudouridine is known as the most abundant and earliest modified ribonucleoside among more than 100 types of RNA post-transcriptional modifications that have been discovered so far [1–3]. Because of its prevalence in cellular RNAs, it has also been considered as the fifth ribonucleoside [1]. The properties, chemical structures and distribution of a pseudouridine are quiet different from its parental base (i.e., uridine) [4, 5]. Compared to uridine, the chemical conformation of a pseudouridine allows the formation of an extra hydrogen bond at its non-Waston-Crick edge [6]. This fact indicates that a pseudouridine can form a more stable base stacking conformation [7], which is believed to play an important role in stabilizing RNA structure [8]. In ribosomal RNAs (rRNAs), it has been found that pseudouridines are required for the maintenance of ribosome-ligand interactions and translational fidelity [9]. In addition, pseudouridines in transfer RNAs (tRNAs) can reduce the conformational mobility of the structural elements around the modified sites and thus affect the amino acid transfer efficiency [10]. Such a stabilization function of pseudouridines in anticodon stem-loop of tRNA^*Lys*,3^ has also been validated by NMR spectroscopy [11]. Moreover, pseudouridines in spliceosomal RNAs can be involved in the RNP assembling and pre-mRNA splicing processes [12]. The above findings indicate that most likely pseudouridines are tightly related to RNA structure stabilization, translation process and RNA stability. However, the underlying mechanisms of pseudouridines’ involvement in the aforementioned processes still remain poorly understood.

The conversion from a uridine to a pseudouridine is catalysed by pseudouridine synthases (PUSs) through two distinct processes, including RNA-dependent and RNA-independent operations [13]. The RNA-dependent pseudouridylation process depends on the box H/ACA ribonucleoproteins (RBPs), which consist of a small box H/ACA RNA and four core proteins, including centromere-binding factor 5 (Cbf5; also known as dyskerin in mammals), non-histone protein 2 (Nhp2), glycine-arginine-rich protein 1 (Gar1) and nucleolar protein 10 (Nop10), to form a pseudouridylation pocket for substrate recognition and catalytic activity [14, 15]. In the RNA-independent pseudouridylation mechanism, a single synthase protein, such as PUS7, is responsible for both substrate recognition and pseudouridylation catalysis [13]. Although notable progress has been made in recent years in studying pseudouridylation in various types of RNAs, the detailed underlying mechanisms of pseudouridylation are still not completely deciphered. So far, about 13 types of PUSs in human have been identified [16], and generally it is difficult to unveil the consensus catalytic laws of pseudouridylation. Moreover, it has been shown that pseudouridylation in RNAs is highly dynamic and inducible [13], which makes it even harder to characterize the properties of pseudouridylation. On the other hand, RNA modification is mostly a sequence pattern recognition process, as it is heavily dependent on the sequence binding preferences of catalytic proteins [17]. From this point of view, it is reasonable to speculate that RNA pseudouridylation is determined by the sequence contexts of the sites being modified.

To characterize the properties of pseudouridines computationally, we need develop efficient methods to accurately identify pseudouridines at single-base resolution and obtain a transcriptome-wide map of pseudouridines. Traditional pseudouridine detection methods are mainly based on the N_3_-CMC labeling and gel electrophoresis experiments [18], which are often labourious and time-consuming. Recently, several high-throughput profiling techniques, including Pseudo-seq, Ψ-seq, PSI-seq, and CeU-seq, have been proposed to map RNA pseudouridine sites to reference transcriptomes [19–22]. These high-throughput experiments typically combine CMC derivatives with next generation sequencing or deep sequencing techniques to detect pseudouridine sites on a transcriptome-wide scale. However, these experiments are generally costly and often require tremendous time and effort in deriving the positions of pseudouridine sites. On the other hand, although plenty of pseudouridine sites in small cellular RNAs have been identified, their sequence contexts have not been fully exploited to gain better insights into understanding their functions. In addition, although recent high-throughput sequencing techniques, such as Ψ-seq and CeU-seq, have been able to identify large-scale pseudouridylation sites in mRNAs, they may still miss numerous modification sites due to the limitations of current experimental methods (e.g., the incompleteness of CMC-labelling or the read mappability issue). Development of efficient machine learning approaches to capture the intrinsic sequence patterns around pseudouridylation sites will enable us to better understand the sequence contexts of pseudouridylation and predict novel pseudouridine sites that are missed by current experimental methods. In addition, by fully exploiting the underlying sequence patterns of pseudouridylation, computational prediction methods may also provide novel predictions of potential pseudouridines in new cell types or species. Moreover, the computational prediction of transcriptome-wide pseudouridylation sites and characterization of their sequence contexts may provide important hints in understanding the functional roles of pseudouridylation in RNA regulation. Although several computational approaches and web servers, such as PPUS [23] and iRNA-PseU [24], have been developed to predict novel pseudouridine sites, they either can only be applied to predict PUS specific sites or only use the handcrafted features derived from the chemical properties of nucleotides.

Recently, deep learning techniques, especially convolutional neural networks (CNNs), have been widely used in genomic data analysis for extracting accurate sequence features [25–29]. CNNs were first developed for handwriting recognition and face identification [30], and have become one of the most famous and powerful learning models in the field of computer vision, speech recognition and natural language processing [31–33]. Despite its powerful predictive capacity, it remains unknown whether a CNN model can be used to effectively capture the contextual sequence features of pseudouridylation and accurately predict new pseudouridine sites.

In this study, we have developed a computational framework, called PULSE (<u>P</u>seudo<u>U</u>ridy<u>L</u>ation <u>S</u>ites <u>E</u>stimator), to predict novel pseudouridine sites from large-scale profiling data of pseudouridines based on the sequence contexts of target sites. To our knowledge, our study is the *first* machine learning based attempt to characterize the contextual sequence features of pseudouridylation by fully exploiting the currently available large-scale profiling data of pseudouridines. PULSE employs a CNN model, which contains a number of alternatively-stacked convolution and pooling layers responsible for local feature extraction from the input contextual sequences and several fully-connected layers responsible for feature integration and estimation of the pseudouridylation potential of a candidate site. Tests on both human and mouse data have demonstrated that PULSE can achieve high prediction accuracy and outperform a conventional machine learning approach called gkmSVM [34]. In addition, the sequence features captured by PULSE are not only consistent with the recognized motifs of known pseudouridylation synthases, but also match the binding patterns of several nucleotide-binding proteins, which may provide useful hints for discovering new potential pseudouridylation synthases.

## Results

### The PULSE framework

We have developed a convolutional neural network (CNN) based framework, called PULSE (<u>P</u>seudo<u>U</u>ridy<u>L</u>ation <u>S</u>ites <u>E</u>stimator), to predict new pseudouridine sites from large-scale pseudouridylation profiling data (Fig. 1a). To encode the contextual sequence features of a potential pseudouridine site of interest, we first extend the target site both upstream and downstream by 50 nucleotides (nts) and then use a simple four-dimensional binary vector to encode each nucleotide (Fig. 1a; Methods). Then, the encoded matrix of an input contextual sequence is fed into a particularly-designed CNN model to capture the latent features of the potential sequence determinants of the pseudouridine site. In particular, our CNN model consists of two convolution layers and two pooling layers, which are alternately stacked and then followed by a fully-connected multilayer network (Fig. 1b; Methods). Generally speaking, the convolution-pooling layers are responsible for local feature extraction, while the fully-connected layers are mainly used for feature integration and final classification [27]. In particular, the convolution kernels from the convolution layers scan the input matrix that encodes the input sequence profiles and capture intrinsic hidden features about the local contextual patterns of the target site. The last fully-connected layer (also called the output layer) employs a softmax function to perform the classification task.

**Figure 1.**
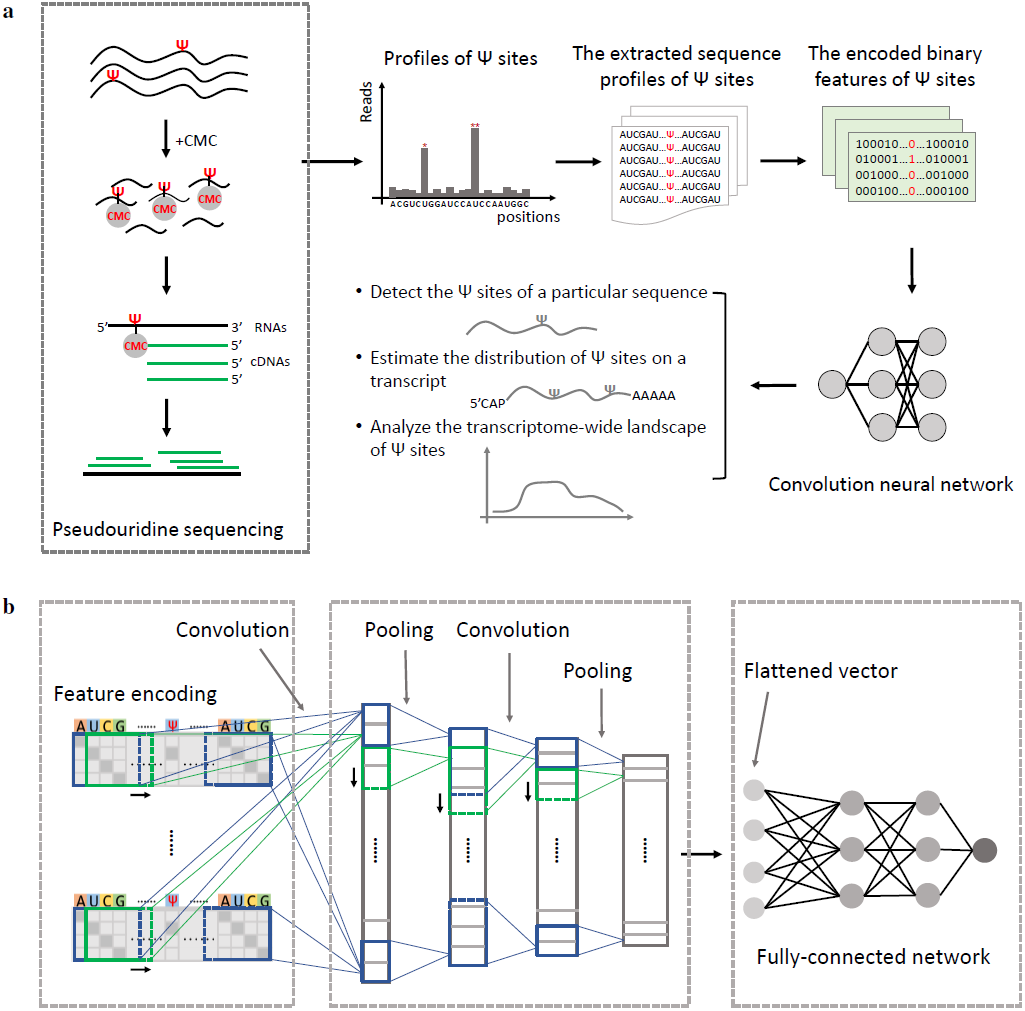
Overview of the PULSE pipeline. **(a)** The schematic flow diagram of PULSE. The pseudouridine (Ψ) sites can be experimentally profiled by high-throughput sequencing techniques, such as Ceu-Seq, Pseudo-Seq, Ψ-seq and PSI-Seq. PULSE first extracts the sequence profiles of a potential Ψ site (i.e., the region within 50nts upstream and downstream of the target site of interest) and encodes them into binary features, which are then fed as input data to a particularly designed convolutional neural network (CNN) model. After parameter learning, the trained model is used for downstream analysis, such as detecting the Ψ sites of a given RNA sequence, estimating the distribution of Ψ sites on a transcript and elucidating the transcriptome-wide landscape of Ψ sites. **(b)** The convolution neural network architecture used in PULSE. Two convolution layers and two pooling layers are first alternately stacked and used for feature detection, and then a fully-connected network with three hidden layers is added for global feature evaluation and Ψ potential estimation. Given a uridine site of interest and its sequence context, PULSE outputs a Ψ potential score which basically represents the likelihood of pseudouridylation for this target site.

We used grid search with a cross-validation procedure [35] to calibrate the hyperparameters of our CNN model (see Methods). In the final optimal setting of the hyperparameters for the first and the second convolution layers, the numbers of local motif detectors were set to 64 and 32, respectively, and the sizes of convolution kernels were set to 4 × 8 and 1 × 8, respectively. The sizes of max-pooling kernels were set to 1 × 2 for both pooling layers. The number of hidden layers (before the output layer) and the number of units in each hidden layer are set to 2 and 64 in the fully-connected network, respectively. For a given sequence l, the overall information flow of PULSE can be abstracted into the following formulas:

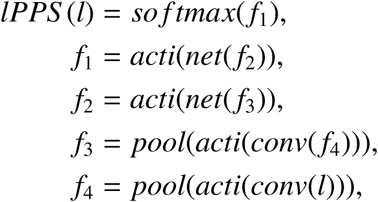
where *lPPS (l)* represents the final prediction score of the target site, and *conv(), acti(), pool()* and *net()* stand for the convolution, neuron activation, pooling and full-connection operations, respectively. In our framework, the length of the input sequence *l* is set to 101, as we extend the target site both upstream and downstream by 50 nts.

### Validating PULSE

To evaluate the prediction performance of PULSE, we applied a 10-fold cross-validation procedure on both human and mouse data (see Methods). In our training process, the pseudouridine sites identified by high-throughput profiling experiments and the corresponding flanking regions of size 50 nts on both sides of individual pseudouridine sites were considered positive samples, while uridine sites with flanking windows of 50 nts on both sides that are the closest to some pseudouridine sites and do not have any overlap with the positive samples were considered as the negative samples. We trained PULSE on human and mouse datasets separately, resulting in two trained models called hPULSE and mPULSE, respectively. We also compared the prediction performance of PULSE to that of a baseline approach, called gkmSVM, which is a classical SVM-based classifier based on gapped k-mers [34]. Our 10-fold cross-validation tests showed that both hPULSE and mPULSE can achieve high prediction accuracy, with the area under ROC curve (AUC) scores 0.86 and 0.84, respectively, which were significantly better than those of gkmSVM (Figs. 2a-2b and Figs. S1a-S1b).

**Figure 2.**
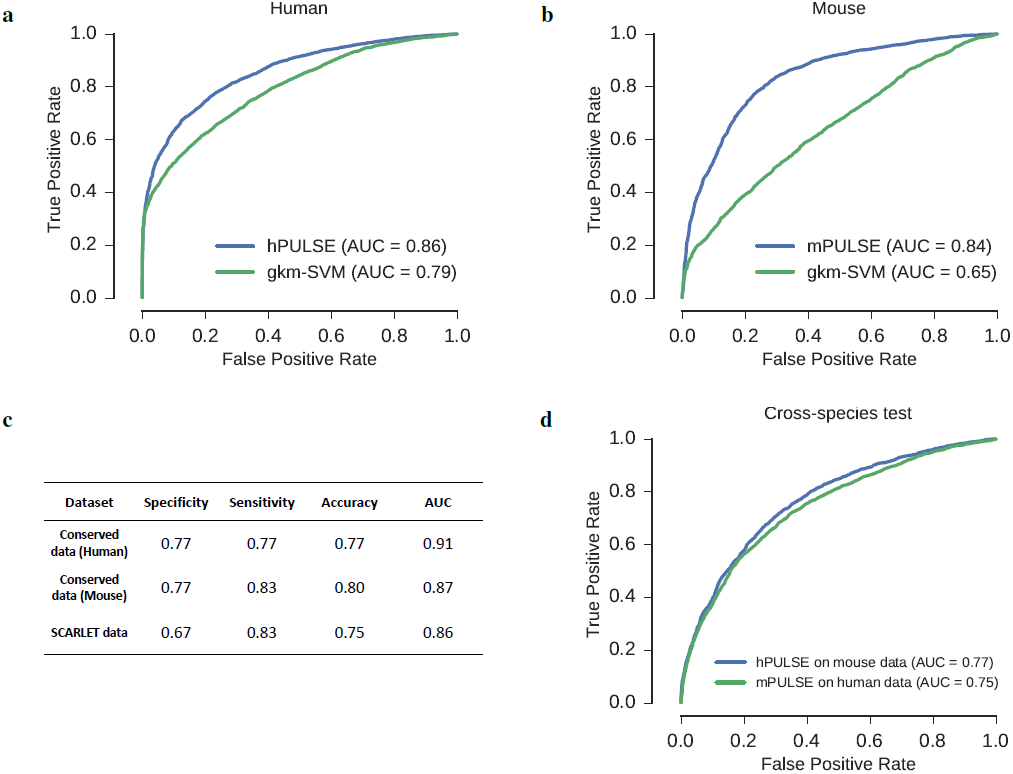
Performance evaluation of PULSE. **(a-b)** The ROC (receiver operating characteristic) curves and the corresponding AUC (area under the ROC curve) scores reported in 10-fold cross-validation for human and mouse, respectively. The terms ‘hPULSE’ and ‘mPULSE’ represent the PULSE models trained based on human and mouse datasets, respectively. gkm-SVM, which is a classical model for sequence classification based on the gapped k-mers [34], was used as a baseline method for comparison. **(c)** The validation results on the conserved pseudouridine sites across human and mouse [22] and a set of the pseudouridine sites in human [22] that were identified by the SCARLET (site-specific cleavage and radioactive-labeling followed by ligation-assisted extraction and thin-layer chromatography) technique [36]. During the evaluation process, for both hPULSE and mPULSE models, the overlapping pseudouridine sites were removed from training data. **(d)** The results on the cross-species test between human and mouse. Green and red curves represent the test results on the hPULSE model (which was trained from human data) over mouse data and the mPULSE model (which was trained from mouse data) over human data, respectively.

To further validate the prediction accuracy of PULSE, we also tested it on three small but relatively more reliable datasets of pseudouridine sites, including a dataset of high fidelity pseudouridine sites that were identified by the SCARLET (site-specific cleavage and radioactive-labeling followed by ligation-assisted extraction and thin-layer chromatography) technique [36], and two datasets of the conserved pseudouridine sites for both human and mouse [22]. In these new validation tests, all overlapping sites were removed from the training data. Our validation tests on these three additional datasets of high-fidelity pseudouridine sites showed that PULSE can still achieve high classification accuracy, with the AUC scores between 0.86-0.91 (Fig. 2c).

Previous studies showed that the pseudouridylation profiles of transcriptome across human and mouse were conserved to some extent despite the possible difference in the underlying mechanisms of pseudouridylation [22]. Thus, we speculated that the conservation of pseudouridine sites between human and mouse can also be reflected by the prediction results of PULSE. To test this speculation, we performed a cross-species test between human and mouse datasets, that is, we used the PULSE model trained from human (mouse) pseudouridylation profiles to test the mouse (human, respectively) data. As expected, such a cross-species test demonstrated a strong conservation relationship between human and mouse in pseudouridyaltion (Fig. 2d and Fig. S1c). In addition, this cross-species validation test also implied an impressive generalization capacity of PULSE in predicting new pseudouridine sites.

In summary, the above validation tests implied that PULSE can effectively recognize the underlying latent features of the sequence contexts of pseudouridylation and thus yield accurate predictions of pseudouridine sites.

### The motifs of pseudouridylation captured by PULSE

We further analysed and visualized the motifs of the sequence contexts of pseudouridylation captured by the filters employed in the first convolution layer of PULSE, using the same strategy as in the previous studies [25, 27, 37]. In particular, we focused on high confident motifs that covered more than 1% (about 50) of the positive samples in the training data. As a consequence, we obtained 300 and 272 sequence patterns identified by PULSE for human and mouse, respectively (see Methods).

As expected, we found that the previously-known sequence recognition motifs of pseudouridine synthases PUS4 and PUS7 [22, 38–40], i.e., ‘GUΨCNA’ and ‘UGΨAG’, appeared repetitively in the sequence patterns identified by the filters of our CNN model for both human and mouse (Figs. 3a-3b). We also used the RSAT tool [41] to cluster all these sequence patterns identified by PULSE and obtained 70 and 69 clusters for human and mouse, respectively (Fig. S2; Methods). These motif clusters may provide useful insight into understanding the underlying mechanisms of pseudouridylation. Intriguingly, several novel motifs also appeared repetitively in our visualization results. We hypothesize that these motifs may correspond to other pseudouridine synthases or recognition proteins. Thus, we mapped our discovered motifs to the known binding motifs of RNA binding proteins (RBPs) from the CIS-BP database [42] and transcription factors (TFs) from the HOCOMOCO database [43] (see Methods). As a result, several of the newly discovered sequence motifs of pseudouridylation significantly match the known binding motifs of both human and mouse (*P* < 10^−4^; Figs. 3c-3d). We found that these matching motifs were highly related to important RBPs and TFs, e.g., PCBP1, an RBP involved in the regulation of alternative splicing [44], and FOXO3, a TF acting as a trigger of apoptosis [45]. These discovered novel motifs that matched the known binding sequence patterns of RBPs implied that the RBPs may play important functional roles in the pseudouridylation process, which thus may also provide new candidate molecules of pseudouridine synthases for further experimental studies. Since previous studies showed that RNAs can also be co-transcriptionally modified [46], the TFs with matching sequence motifs may be related to the triggers of pseudouridylation during RNA transcription.

**Figure 3.**
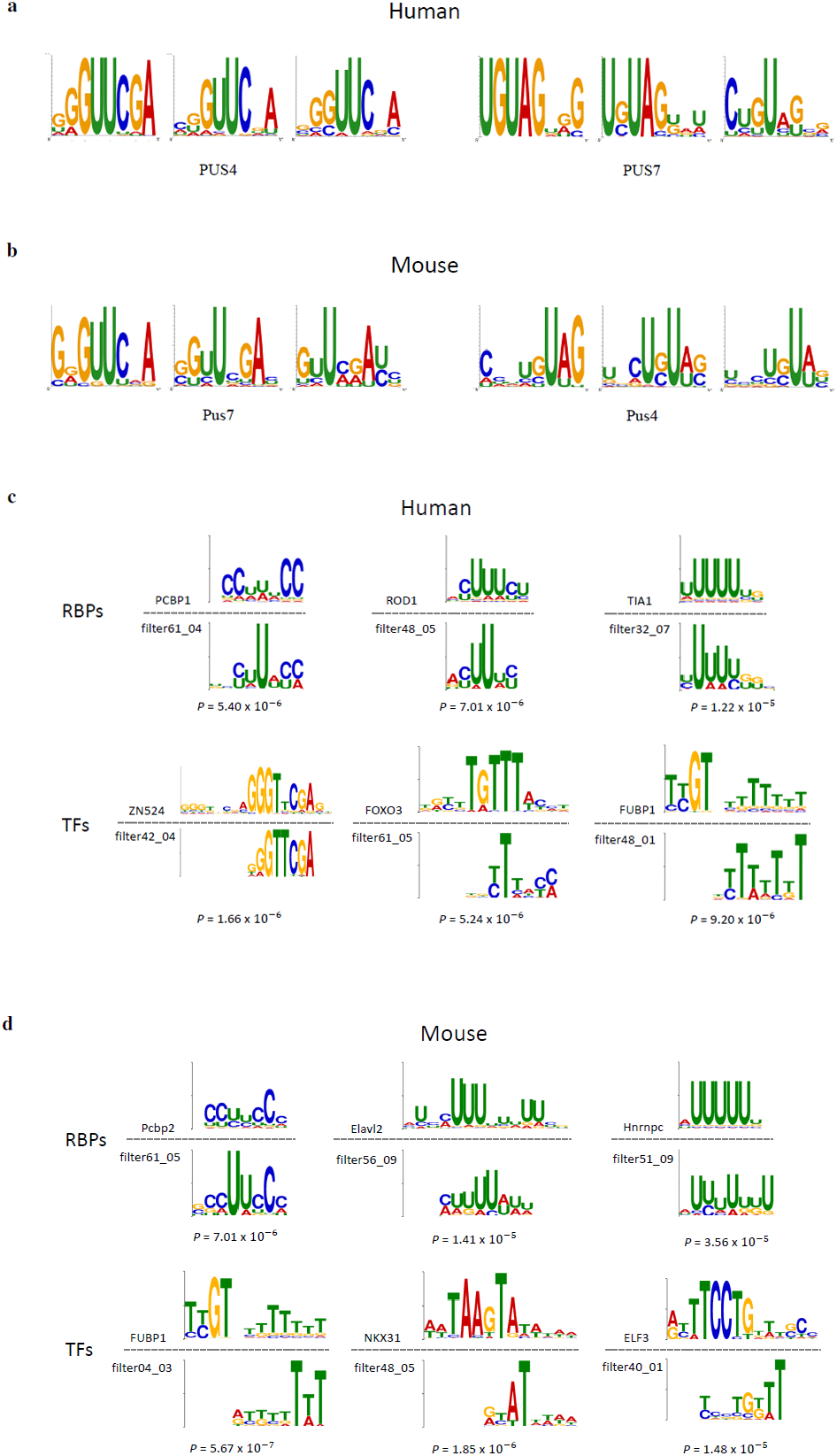
Examples of the sequence motifs of pseudouridylation identified by PULSE. **(a-b)** The sequence motifs of pseudouridylation detected by PULSE corresponding to the known motifs of Ψ sythases PUS4 (‘GUΨCNA’) and PUS7 (‘UGΨAG’) for human and mouse, respectively. **(c-d)** The comparisons between the motifs of the sequence contexts of pseudouridylation identified by PULSE and the closest matched motifs of RNA binding proteins (RBPs) and transcription factors (TFs) for human and mouse, respectively. Top and bottom show the known binding motifs of RBPs or TFs and the contextual sequence features of pseudouridylation identified by PULSE (the filter IDs are also showed), respectively.

### Transcriptome-level characteristics of the detected pseudouridine sites

The trained PULSE model allow us to unveil the landscape of pseudouridylation across the either (human and mouse) transcriptome. As mentioned above, each uridine in a transcript can be characterized by a local pseudouridylation potential score (1PPS) derived from PULSE based on its corresponding sequence context. Basically, this lPPS value measures the probability that a uridine can be converted into a pseudouridine. Based on the distribution of uridines on a transcript and the corresponding lPPS profiles, we derived a new metric, called the transcript pseudouridylation potential score (tPPS; see Methods), to estimate the overall pseudouridylation level of this transcript. Based on the 1PPS and tPPS profiles derived from PULSE, we are able to study the nucleotide- and transcript-level landscapes of pseudouridylation, respectively.

Modifications in different regions of mRNAs may play distinct roles in determining their fates. For example, *N*^6^-methyladenosine (m6A) in the coding sequence (CDS) and the 3’UTR of an mRNA can modulate its translation efficiency [47], while m6A in specific intronic regions can influence the splicing results [48, 49]. To examine the transcriptome-wide distribution of pseudouridylation across different genomic regions, we compared the percentages of pseudouridylation predicted by PULSE among different types of regions, including 5’UTRs, CDSs and 3’UTRs. Our comparison showed that pseudouridines appear primarily in the CDS regions (about 50%) and the 3’UTRs (about 40%) of both human and mouse mRNAs (Fig. 4a), which was consistent with the previous reports [22, 40]. As the 3’UTRs of mRNAs are tightly associated with RNA stability and translational control [50, 51] and the CDS regions contain the core genomic contents that are translated into proteins, it is reasonable to hypothesize that the pseudouridylation activities in the 3’UTRs and CDS regions are involved in RNA stability modulation and translational regulation.

**Figure 4.**
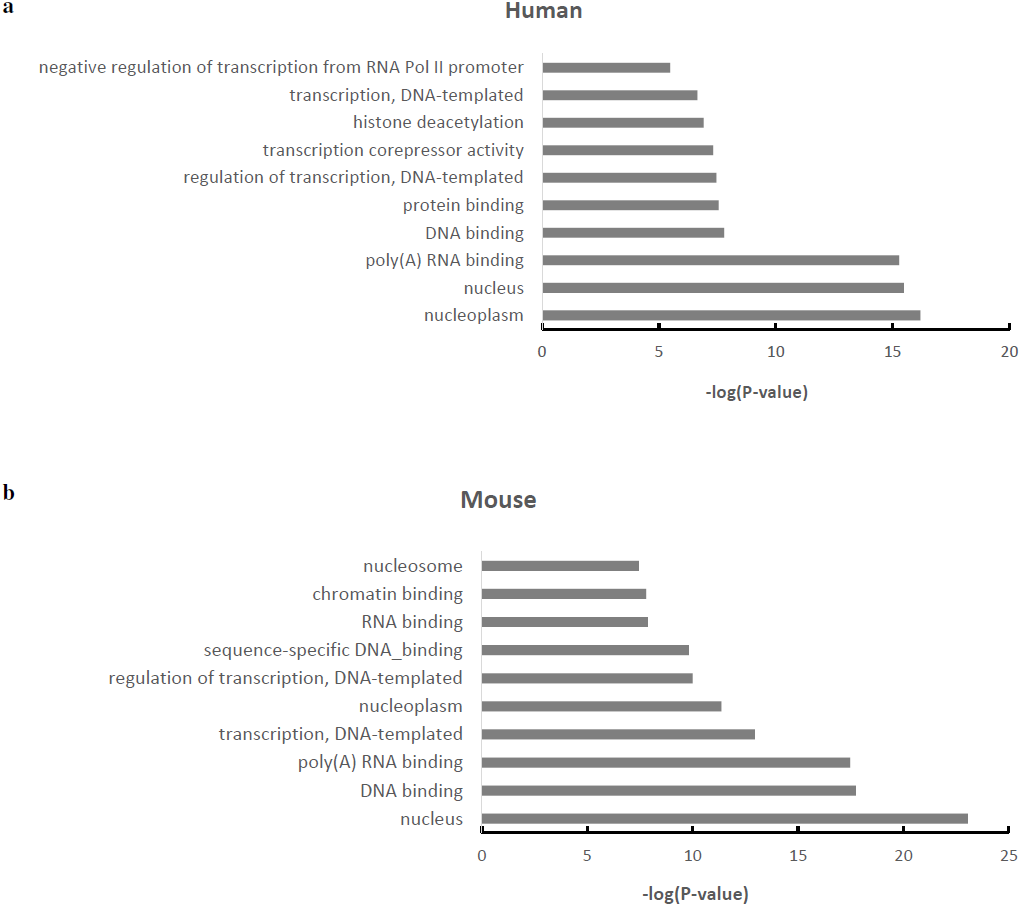
The gene ontology (GO) enrichment analysis of mRNAs with high tPPS values (top 500) carried out by DAVID for human **(a)** and mouse **(b)**, respectively.

### GO enrichment analysis

Gene ontology (GO) enrichment analysis can highlight the consensus functions of a particular gene set, which may help reveal the latent features of the genes in the set [52]. To further elucidate the potential functional roles of pseudouridylation in mRNAs, we performed GO enrichment analysis for the top 500 genes with the highest transcript-level pseudouridylation potential scores (tPPSs) for both human and mouse. We found that genes associated with high tPPS values predicted by PULSE are mainly distributed in the nucleus and contribute to DNA or RNA binding (Figs. 8a-8b). This phenomenon implied that pseudouridines in mRNAs may also serve as molecular glues between nucleotide-binding proteins and RNAs by increasing their interaction strength and forming more stable complex conformations. This hypothesis is in line with the previous results about the potential functions of pseudouridylation in RNA secondary structure and translational regulation [5, 7, 53]. Of course, we will need extensive experimental studies and more experimental data to further validate the hypothesis in the future.

## Discussion

Although numerous studies have revealed that pseudouridines in small cellular RNAs are important for maintaining their structures [5, 7, 11], the functional roles of pseudouridylation in mRNA translation and its contextual sequence features still remain largely unclear. Several high-throughput experimental techniques of characterizing pseudouridines across transcriptome have been developed recently [19–22], detailed transcriptome-wide landscape, functions and molecular mechanisms of pseudouridines in post-transcriptional regulation are still to be revealed. In this study, we developed an effective convolution neural network (CNN) model to detect pseudouridine sites, based on which we further analysed the landscape of pseudouridines across the human and mouse transcriptomes. Our model can not only capture the known motifs of pseudouridylation that were consistent with previous studies, but also reveal novel motifs that may help uncover potential new pseudouridylation synthases. Moreover, the GO enrichment analysis of genes with high pseudouridylation scores predicted by our model may provide a wide vision on the potential functions of pseudouridylation.

RNA pseudouridylation is obviously crucial to RNA regulation simply by its prevalence in transcriptome and its high conservation across different species. Therefore, a comprehensive understanding of RNA pseudouridylation will be conducive to the consummate studies of RNA modifications and RNA epigenetics. The studies of RNA pseudouridylattion especially in mRNAs may help understand its functional roles in post-transcriptional regulation. On the other hand, currently, numerous aspects of RNA pseudouridylation including its functions in RNA stability, alternative splicing, RNA localization and protein binding, still remain largely unknown. Our model in this study may provide a new avenue for further exploration in the aforementioned fields. In addition, our proposed deep learning model may provide a new paradigm for analysing other types of RNA modifications, such as 5-methylcytidine (m5C) and N1-methyladenosine (m1A) [54–56], which can also help decipher the RNA modification code and understand their regulatory functions in gene activities.

## Methods

### Data collection and preprocessing

The pseudouridine modification site data were downloaded from the RMBase database [57] which includes the high-throughput profiling data of pseudouridines collected from three recent experimental strudies [19, 20, 22]. All the labelled pseudouridine sites were separated into a human dataset and a mouse dataset. In addition, the overlap between human and mouse datasets which represents the conserved pseudouridine sites was considered a relatively reliable dataset for further model validation. Moreover, the pseudouridylation sites identified by SCARLET curated from [22] were also used for assessing the prediction accuracy of PULSE. All of the above modification sites were mapped to the reference genome (human: hg19; mouse: mm10) and those that cannot be mapped to thymines were discarded. A sequence of length 101 that covers the pseudouridine site and has a 50-nt window flanking on its both sides was labelled as a positive sample, while the sequence of the same length that is centered at a thymine that is closest to a corresponding Ψ site and does not have any overlap with any positive sample was labelled as a negative sample. In the end, we collected 7720 and 6166 samples in total for human and mouse, respectively.

### Feature encoding of the sequence profiles

We use the following scheme to encode individual nucleotides in a given input sequence *S*,

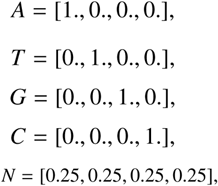
where ‘N’ stands for any nucleotide. Then a given sequence can be encoded into a matrix. For example, sequence *S* = ‘ATGCAN’ can be encoded as

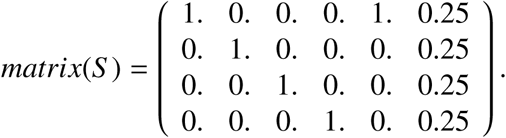

### Model construction and training

The core of PULSE is a convolutional neural network (CNN), which has become a powerful deep learning model in various data science fields, such as computer vision [31], speech recognition [32] and natural language processing [33]. Recently, CNNs have also been successfully applied to capture the latent features of genomic data, e.g., estimation of sequence specificities of DNA or RNA binding proteins [27], prediction of functional effects of non-coding variants [28] and characterization of splicing codes [29]. A CNN model typically consists of several alternately-stacked convolution and pooling layers which basically act as motif detectors. During convolution, the input matrices of dimension *L* × 4 (where *L* stands for the length of the input sequences) are first cross-correlated with several convolution filters and then the convolved outputs are rectified by a particular activation function (e.g., sigmoid or ReLU activation). In the pooling stage, the pooling operators are applied to the previous convolution and activation results for further motif extraction. After that, the pooled results are flattened to a vector which is then fed to a fully-connected neural network for final classification. In particular, our computational pipeline PULSE is composed of two alternate convolution and pooling layers, which are then stacked with a fully-connected neural network consisting of three hidden layers (Fig. 1b).

More specifically, given an RNA sequence of length *L* and the corresponding feature-encoding matrix *S*, the convolution result *X* = *conv*(*S*) is an (*L* – *m_c_* + 1) × *n* array, and can be written as

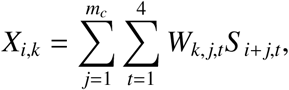
where *n* and *m_c_* stand for the number and the size of convolution operators, respectively, 1 ≤ *i* ≤ *L* − *m_c_* + 1,1 ≤ *k* ≤ *n*, and *W* represents the weight matrix of the *n* convolution operators, i.e., *W_k,j,l_* represents the value at the *j-th* row and the *l-th* column of the *k-th* kernel. In the rectification stage, the parametric rectified linear activation function (PReLU) [58] is applied to the previous convolved result *X*, that is,

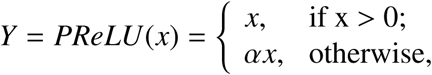
where *α* ≥ 0 stands for the slope of the negative part which can be learned in the model training process.

In the pooling stage, the dimension of *Y* is reduced by a max-pooling operator of size *m_p_*. Suppose that the pooling result is defined as *Z = max_pool(Y)*. Then we have

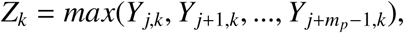
where 1 ≤ *k* ≤ *n* stands for the index of the pooling operator and *j* stands for the start position of pooling.

Next, the output matrix *Z* of max-pooling is PReLU rectified, flattened to a high-dimensional vector and then fed to a fully-connected neural network, which can be formulated as *Z_net_* = *W_net_ · PReLU*(Z), where *W_net_* represents the weight matrix of the fully-connected network. For final classification, a softmax layer is employed in the last layer of the fully-connected network, that is,

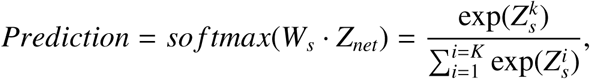
where *W_s_* stands for the weight matrix of the softmax layer, *Z_net_* stands for the output of the proceeding layer in the fully-connected network, and *K* stands for the length of *Z_s_*.

We applied a grid search like procedure to scan the possible ranges and calibrate those hyperparameters related to the network structure, such as the size and the number of convolution operators. In the final setting of PULSE, the sizes of the first and the second convolution layers were set to 4 × 8 and 1 × 8, respectively, and the sizes of both pooling layers were set to 1 × 2. The numbers of convolution operators in the first and second layers were set to 64 and 32, respectively, and the numbers of units in the hidden layers of the fully-connected neural network were set to 64-64-1. We then applied a 10-fold cross-validation procedure to train PULSE (i.e., estimating the weighting parameters in the model) and then evaluate its prediction performance. We first held out about 1/10 of samples as an independent validation dataset to choose the optimal setting of those hyperparameters related to the training process, such as the learning rate and the number of training epoches. We then split the remaining data into 10 parts, and in each fold, one part was used for testing and the others for training. Next, the average values of hyperparameters, e.g., the number of training epoches, over 10 folds were chosen. After that, the previous validation data were merged and the 10-fold cross-validation procedure was performed on all data again to train the model and evaluate its prediction performance based on the previously determined hyperparameters. The human and mouse datasets were used independently to train two separate PULSE models, unless specifically described.

PULSE is implemented based on the Keras library [59] in python. Back propagation is applied in the training process for efficiently updating the parameters [60]. In addition, several optimization techniques, including stochastic gradient descent (SGD), dropout, batch normalization, early stopping and momentum [61–63] are used to improve the training process (e.g., reducing the likelihood of overfitting).

### Motif visualization and analysis

We applied the filters embedded in the first convolution layer of our CNN model to generate the sequence motifs of pseudouridylation, using the same strategy as in [25, 27, 37]. More specifically, we used a window of the size equal to the length of the filters (i.e., 8) to scan the flanking regions on both sides of a pseudouridine site. During this scanning process, those sequence segments (of length 8) with activation values more than half of the maximum score were output. Then these detected sequence segments were converted into the position weight matrix (PWM) form to generate the corresponding motifs representing the sequence contexts of pseudouridylation. To compare these obtained motifs to those known binding patterns of RNA binding proteins (RBPs) and transcription factors (TFs), we searched over the CIS-BP [42] and HOCOMOCO [43] databases (version 2016 for both) using the Tomtom tool [64], respectively, and then clustered all the motifs using RSAT [41] with default parameter settings. The final sequence motifs were visualized using seq2logo [65].

### Transcriptome-wide detection of Ψ sites

We further applied PULSE to detect potential Ψ sites on each transcript along the genome. All the RNA sequences of human and mouse were downloaded from Ensembl [66] by Biomart [67] under references hg19 and mm10, respectively. For each transcript, every thymine site and the flanking 50-nt regions on its both sides were extracted as the input sequence profile to PULSE (‘N’s were padded if the flanking windows were out of the transcripts). Then PULSE computed the local pseudouridylation potential score (1PPS) for each thymine, which measured its pseudouridylation probability.

### Transcript pseudouridylation potential scores (tPPSs)

To evaluate the pseudouridylation potential of a particular transcript, i.e., the estimation of the overall pseudourdylation level of a complete transcript, we defined a new metric called the transcript pseudouridylation potential score (tPPS). In particular, for a transcript *s*, its tPPS value can be defined as follows:

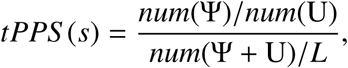
where

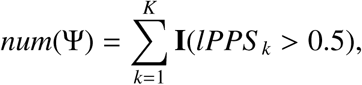

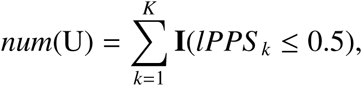
in which *num*(·) represents a count function, *I*(·) represents a binary indicator function, *lPPS_k_* stands for the local pseudouridylation potential score of the *k-th* pseudouridine site in *s, K* represents the total number of thymines in *s*, and *L* stands for the length of *s*. In the above formula, the numerator represents the ratio between pseudouridines and thymines, which thus measures the relative abundance of possible pseudouridines in a transcript. However, this value may bias to those uridine-enriched transcripts. In order to eliminate such bias, the ratio in the numerator is further normalized by the abundance of both thymines and pseudouridines in the transcript.

### GO enrichment analysis

For gene ontology (GO) analysis, we first selected top 500 mRNA transcripts with the highest tPPS values. Then we uploaded the gene names of these 500 transcripts to DAVID [68, 69], and ran the functional annotation clustering module with the default parameters for GO enrichment analysis. During the analysis, the P values were calculated based on the binomial distribution and the Benjamini corrected P values were used for selecting out the final significant GO terms.

**Figure S1.**
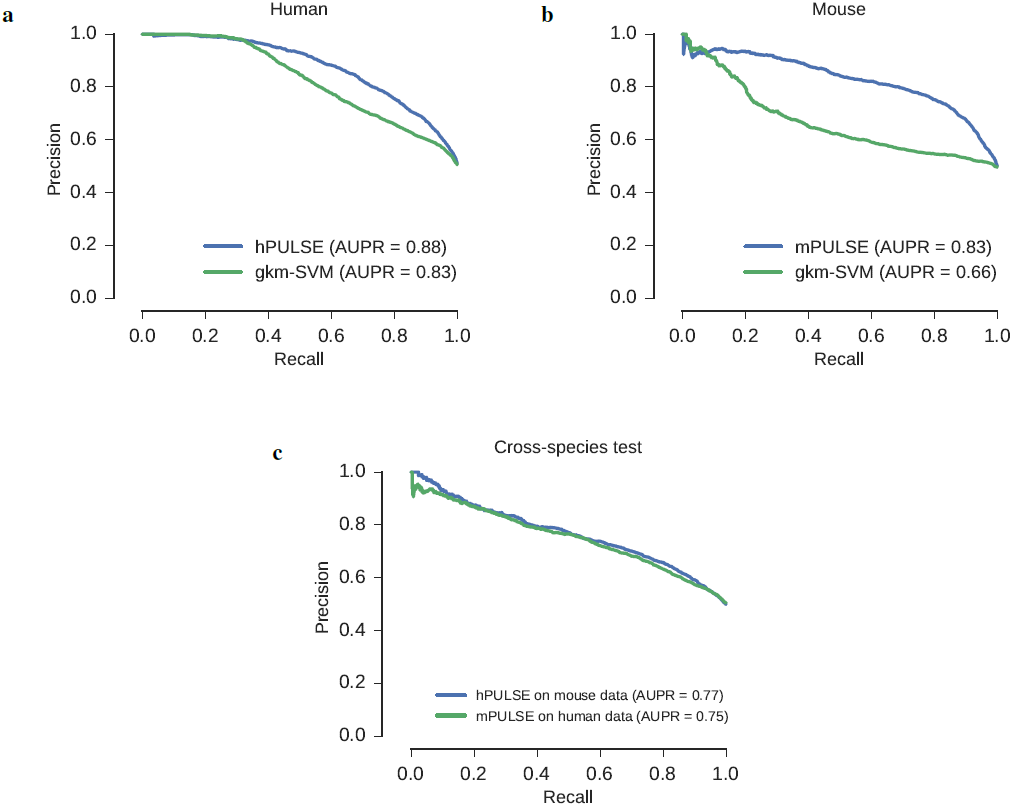
The precision-recall (PR) curves and the corresponding area under the precision-recall curve (AUPR) scores in the cross-validation results. (a-b) Comparisons of the 10-fold cross-validation results between PULSE and the baseline approach gkm-SVM for human and mouse, respectively. (c) The results on cross-species test between human and mouse. The legends are the same as in Fig. 2.

**Figure S2.**
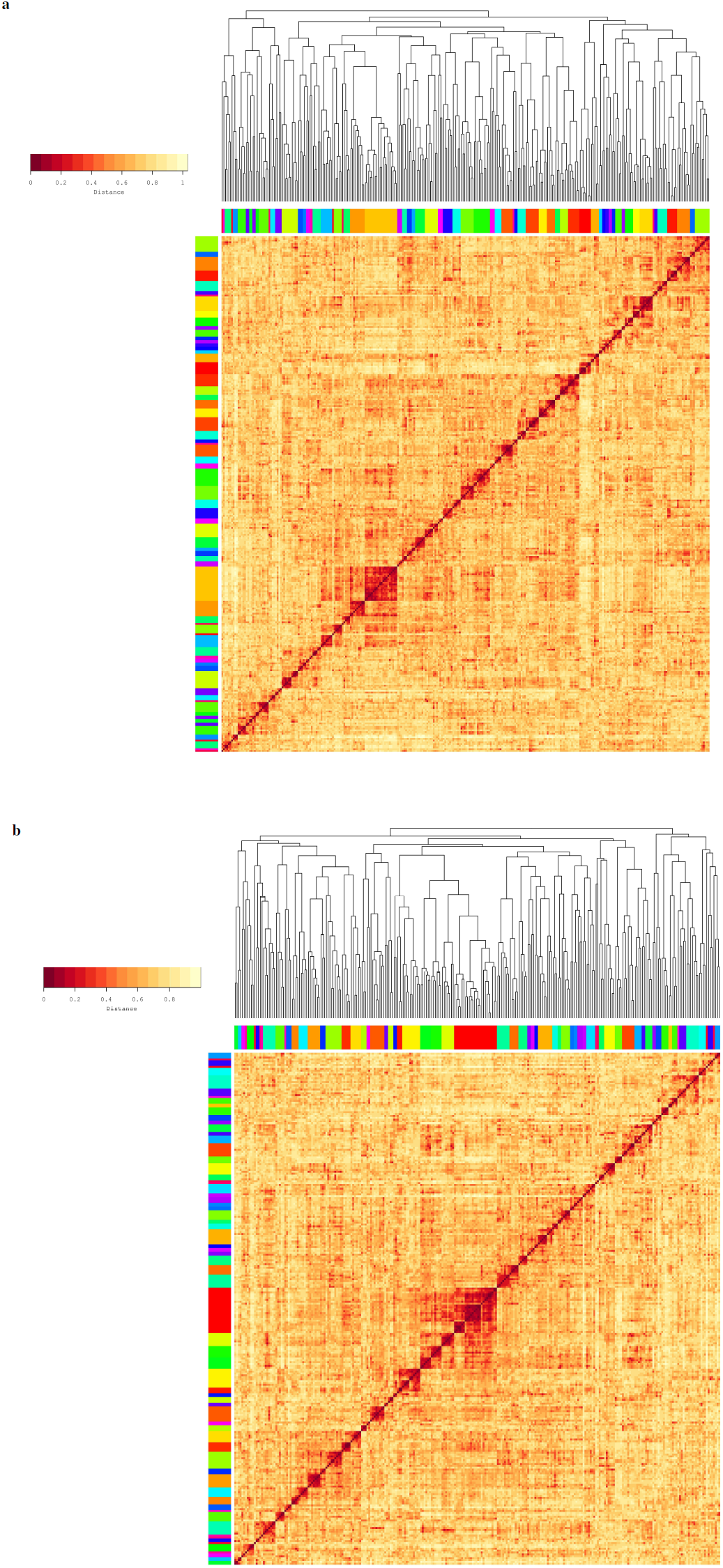
The clustering heatmaps of the sequence motifs of pseudouridylation identified by PULSE for human (**a**) and mouse (**b**). The sequence motifs of human and mouse had 70 and 69 clusters, respectively. The clusters are labelled by different colors.

## Acknowledgments

This work was supported in part by the National Basic Research Program of China Grant2011CBA00300, 2011CBA00301, the National Natural Science Foundation of China Grant 61033001, 61361136003 and 61472205, the US National Science Foundation grants DBI-1262107 and IIS-1646333 and China′s Youth 1000-Talent Program, the Beijing Advanced Innovation Center for Structural Biology.

## Author contributions

X.H.and J.Z. conceived the research project. J.Z. supervised the research project. X.H. preprocessed raw data, designed and implemented PULSE, and carried out model training, validation and data analysis tasks. Y.Z. helped integrate the data. X.H., S.Z., T.J. and J.Z. fine-tuned the model and performed all the statistical analyses. X.H., T.J. and J.Z. wrote the manuscript. All the authors discussed the results and commented on the manuscript.

